# Long-term population decline of a genetically homogenous continental-wide top Arctic predator

**DOI:** 10.1101/2022.04.29.490071

**Authors:** Marianne Gousy-Leblanc, Jean-François Therrien, Thomas Broquet, Delphine Rioux, Nadine Curt-Grand-Gaudin, Nathalie Tissot, Sophie Tissot, Ildiko Szabo, Laurie Wilson, Jack T. Evans, Victoria Bowes, Gilles Gauthier, Karen L. Wiebe, Glenn Yannic, Nicolas Lecomte

## Abstract

Genetic analysis can provide valuable information for conservation programs by unraveling the demographic trajectory of populations, by estimating effective population size, or by inferring genetic differentiation between populations. Here, we investigated the genetic differentiation within the Snowy Owl (*Bubo scandiacus*), a species identified as vulnerable by the IUCN, to (i) quantify connectivity among wintering areas, (ii) to evaluate current genetic diversity and effective population size and (iii) to infer changes in the historical effective population size changes from the last millennia to the recent past. The Snowy Owl, a highly mobile top predator, breeds across the Arctic tundra which is a region especially sensitive to current climate change. Using SNP-based analyses on Snowy Owls sampled across the North American nonbreeding range, we found an absence of genetic differentiation among individuals located up to 4,650 km apart. Our results suggest high genetic intermixing and effective dispersal at the continental scale despite documented philopatry to nonbreeding sites in winter. Reconstructing the population demographic indicated that North American Snowy Owls have been steadily declining since the Last Glacial Maximum ca 20,000 years ago and concurrently with global increases in temperature. Conservation programs should now consider North American Snowy Owls as a single, genetically homogenous continental-wide population which is most likely sensitive to the long-term global warming occurring since the Last Glacial Maximum.

## INTRODUCTION

Natural ecosystems are currently changing at an unpreceded rate as a consequence of global climate change (IPCC 2018, Taylor *et al*. 2020). These changes are particularly apparent in the Arctic, where climate warming is estimated to be three times as fast as the global average (ACIA 2004, Box *et al*. 2019, Meredith *et al*. 2020). Species that live in the Arctic are therefore facing a variety of changes and challenges (Gilg *et al*. 2012), and may respond by shifting their geographic ranges, adjusting behaviour or phenotypes, or adapting to the new local conditions (Parmesan 2006, Brown *et al*. 2016, Kelly 2019). In a scenario where environments are changing so quickly and in complex ways, genetic information may be an efficient tool to inform conservation and management programs (Schwartz *et al*. 2007, Hoban *et al*. 2021a, Hohenlohe *et al*. 2021). First, understanding genetic structure allows one to infer dispersal of individuals and hence connectivity among populations. The genetic structure of populations is also important for wildlife conservation because it provides a baseline of the geographic scale at which programs should be implemented and helps to define populations and management units (Fraser & Bernatchez 2001, Funk *et al*. 2012, Yannic *et al*. 2016). Furthermore, genetic tools provide key information on the current effective population size of populations (i.e. an essential parameter for monitoring vital rate; Hoban et al. 2013, 2021b), are useful when calculating the loss of diversity due to inbreeding or intraspecific hybridization (Yannic *et al*. 2017), and help to predict the ability of populations to persist and adapt to new environmental conditions (Hoffmann & Sgrò 2011). In addition, the long-term demographic trajectory of a species may also be retrieved from genetic data (Cristofari *et al*. 2018, Cole *et al*. 2019, Cleary *et al*. 2021), which can help to understand how species have responded to past major climatic and environmental changes. Thus, understanding, predicting, and managing biodiversity responses to rapid climate change demand a full consideration of the genetic differentiation within a species, an assessment of the species’ evolutionary potential and knowledge of demographic trends.

The Snowy Owl (*Bubo scandiacus*), declining according to the IUCN (BirdLife International 2020), provides a case study of how conservation actions can be informed and refined using genetic information. This owl is a highly mobile top predator which breeds exclusively on the Arctic tundra (Holt et al. 2020). In Canada, Snowy Owls are declining at an estimated −0.03 to - 2.85% rate per year depending on the region (Christmas bird counts; Meehan et al. 2018). Using very different estimation and extrapolation methods, the global Snowy Owl breeding population was first estimated at *ca* 290,000 individuals (Rich *et al*. 2004) and then more recently at *ca* 28,000 mature individuals (Potapov & Sale 2012). Both estimates are, however, probably inaccurate because they are not based on actual continental-wide surveys (Fuller *et al*. 2003, Therrien *et al*. 2014, Holt *et al*. 2020).

Currently, there is little genetic information on Snowy Owl populations. A previous study based on mtDNA concluded there was little phylogeographic differentiation across the geographic range of the species, suggesting one global panmictic population (Marthinsen *et al*. 2009). These authors estimated the global species effective population size (*Ne*) at *ca* 14,000 individuals. This estimate, derived from mtDNA, corresponds to the number of breeding females and does not equate to the whole population size as it is considered by IUCN (BirdLife International 2020). Indeed, not only is *Ne* not a direct proxy for census population size (e.g., Ferchaud et al. 2016), but this estimate disregards the number of breeding males or the variance in reproductive success among individuals (Storz *et al*. 2001, Wang *et al*. 2016). Therefore, further research using genome-wide nuclear data would provide a better understanding of the genetic structure of the population and a more accurate estimate of the current effective population size of Snowy Owls.

Snowy Owls exhibit a diversity of movement strategies throughout the year (McCabe *et al*. 2021). During the non-breeding season, individuals regularly overwinter in the Canadian Prairies and the American Great Plains (Chang & Wiebe 2018), and throughout the entire temperate regions of North America during irruption years (Kerlinger & Lein 1988, Robillard *et al*. 2016, Therrien *et al*. 2017, Holt *et al*. 2020). Conversely, some individuals stay in the Arctic throughout winter (Therrien *et al*. 2011, Robillard *et al*. 2018). Snowy Owls from the central regions of North America appear to follow similar and constant migration routes from breeding to winter areas (Curk et al. 2020; K. L. Wiebe personal communication), and other studies concluded there was some site fidelity to wintering areas by Snowy Owls (Therrien *et al*. 2011, Robillard *et al*. 2018). In contrast, individuals from the eastern part of the North-American continent have non-regular and unpredictable movement patterns (Therrien *et al*. 2014, Robillard *et al*. 2018). In summer, there is evidence that Snowy Owls use breeding sites far apart between consecutive years, e.g. 725 km on average and up to 2, 224 km, for individuals in eastern Canada (Therrien et al. (2014) and 1,088 km for individuals in northern Alaska; Fuller et al. 2003), suggesting a lack of breeding site fidelity for the species (Therrien *et al*. 2014, Doyle *et al*. 2017, Holt *et al*. 2020). The large-scale movements of the species among breeding and nonbreeding seasons suggest there could be widespread connectivity among populations. Thus, because Snowy Owls in different parts of their geographic range exhibit both regular and predictable migratory movements and also large-scale unpredictable movements, our primary objective was to study how such movements affect overall genetic structure of the species at a continental scale.

Using SNP data, we specifically tested whether the large dispersal capacity of the species resulted in a high degree of genetic mixing or whether, on a continental scale within North America, the owls showed sufficient wintering site fidelity and non-random migration patterns to exhibit population structure. In a conservation context, we also attempted to estimate the current effective population size (*Ne*) of the species in North America. Our last objective was to investigate the long-term population trajectory of Snowy Owls in North America since the Last Glacial Maximum (~ 21k years ago) and beyond. Understanding how past shifts in climate may have affected population size might provide insight into the resilience of wildlife resilience to climate change in the near future. Past climatic fluctuations deeply affect long-term (historical) effective population size for Arctic terrestrial (Prost *et al*. 2010, Lorenzen *et al*. 2011, Yannic *et al*. 2014) and marine species (Louis *et al*. 2020, Cleary *et al*. 2021), and this never been explored for Snowy Owls, an important species which structures the Arctic trophic network (Legagneux *et al*. 2012, Gauthier *et al*. 2013).

## METHODS

### Sample Collection

We used feathers (*n*=74), blood (*n*=2) and tissue (liver [*n*=48], muscle [*n*=26]) from Snowy Owls collected via live-trapping or carcass collection (brought to rehabilitation centers, government agencies, and veterinary laboratories). Sample collection covered most of the southern nonbreeding region in North America, including British Columbia (*n*=50), Saskatchewan (*n*=29), Minnesota (*n*=3), Ontario (*n*=1), Wisconsin (*n*=9), Michigan (*n*=16), Maryland (*n*=3), Pennsylvania (*n*=2), New York (*n*=34), Québec (Nunavik; *n*=1), Nova Scotia (*n*=1), and Prince Edward Island (*n*=1; Figure 1, Table 1, Supplementary Material Table S1). Sampling spanned a 4,650-km distance. We collected samples during winter between 2012 and 2018 (except for two samples that were collected in 2007 and 2008). For each individual, we assessed sex and age class (first year or adult) according to morphological and molt measurements (Solheim 2012). The spatial distribution of the samples was slightly clustered into four groups (i.e. wintering areas) according to longitude (see Figure 1). We called these wintering areas: West (W: British Columbia); Central (C: Saskatchewan); Great Lakes areas (GL: Michigan, Wisconsin, Ontario, and Maryland); and East (E), which included all other individuals (including the one from Nunavik; Figure 1).

**Figure 1.**
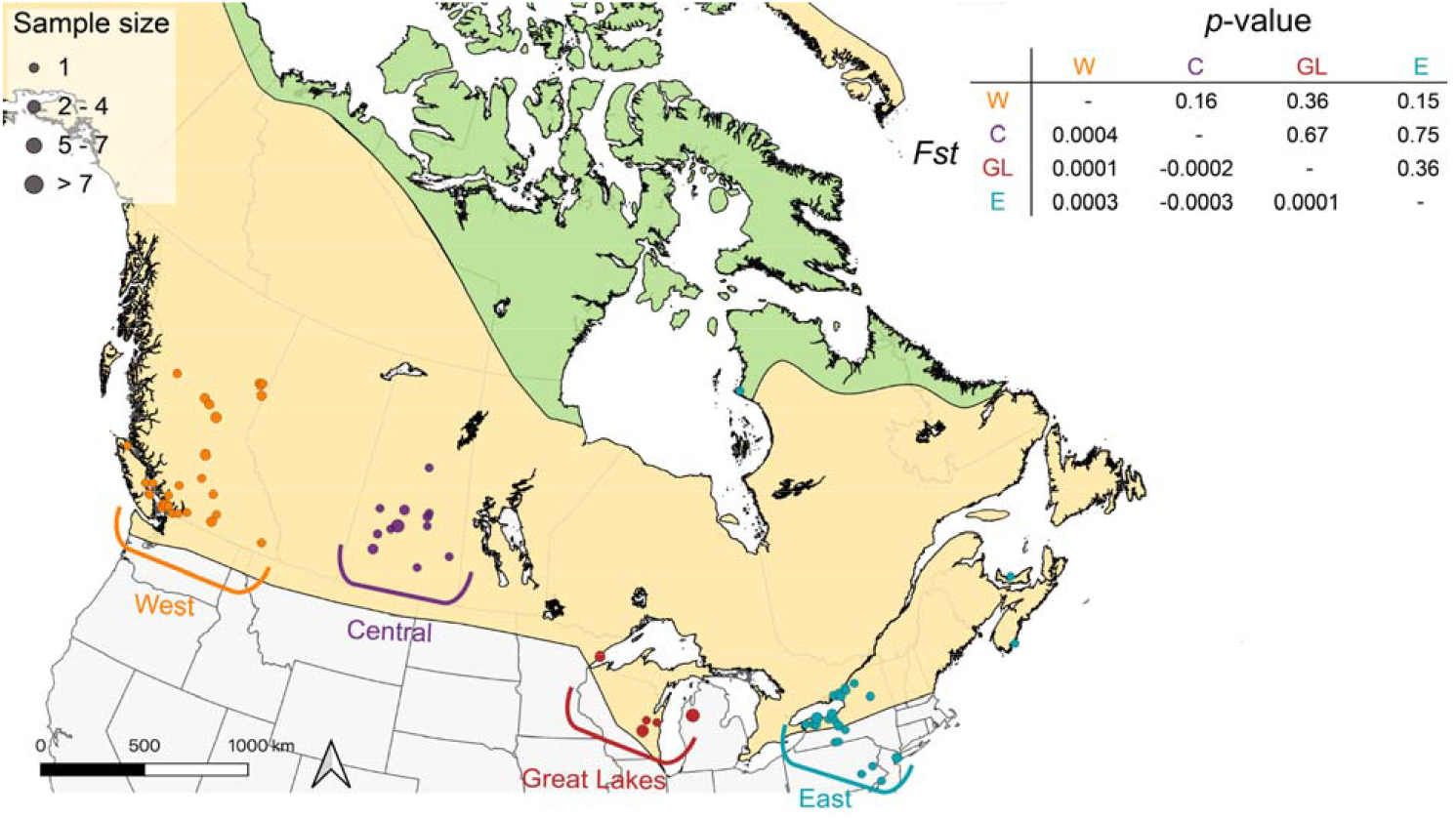
Sampling sites of wintering Snowy Owls (n=150) collected during November-March between 2007 and 2018. Green areas represent the geographic year-round range and yellow areas represent nonbreeding range of the species (BirdLife International 2020). Pairwise *Fst* values among wintering areas (W-West, C-Central, GL-Great Lakes, E-East) and the associated *p*-values are displayed in the legend section.

**Table 1.**
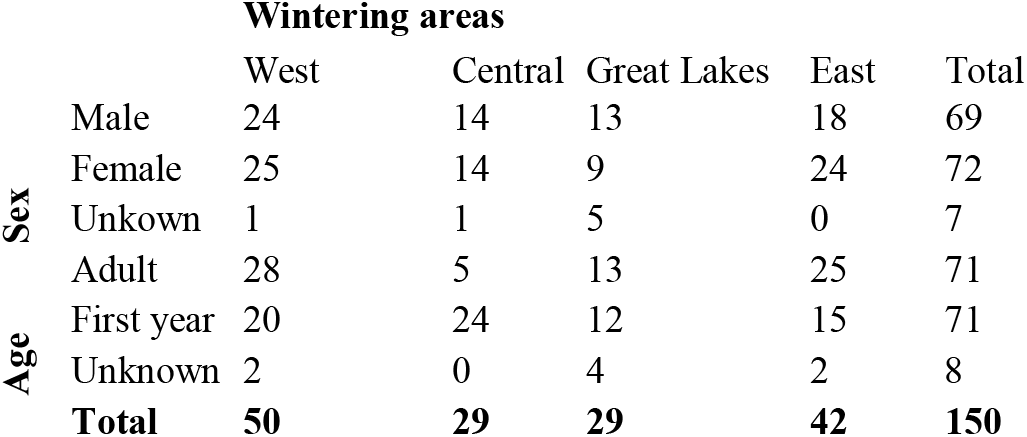
Sampling distribution of Snowy Owls per sex or age for each wintering area.

### DNA Extraction

We used Qiagen DNeasy Blood and Tissue kit (Qiagen, Inc., Valencia, CA, USA) to extract genomic DNA following the manufacturer’s instructions and assessed DNA quality and checked for DNA degradation on agarose gels. We quantified DNA concentration for each sample using the QuantiFluor dsDNA System kit (Promega), using samples with a minimum DNA concentration of 3.20 ng/μl after extraction for subsequent double digested RAD sequencing (ddRADSeq) library preparation.

### ddRADSeq Library Construction and Sequencing

We built four ddRADSeq libraries from the Snowy Owl genomic DNA following a modified version of the protocol in Peterson et al. (2012) and detailled in Gagnon et al. (2019). We randomized the individuals for each wintering area in each library. The Genomic DNA (100 ng) was digested using the enzymes SbfI and MspI and fragments selected between 300 bp and 500 bp using BluePippin size-selection system (Sage Science). To control for library quality and consistency, we replicated 15.2 % of our samples (*n*=28) following the recommendations of Mastretta-Yanes et al. (2015). We included 47 samples per library including replicates and a negative control. The four libraries were then sequenced on two full lanes (two libraries per lane) of paired-end (2 × 125bp) Illumina Hi-Seq 2500 (Fasteris SA, Switzerland).

### ddRADSeq Data Processing

We used Stacks*1.44 (Catchen et al. 2011, 2013) to demultiplex data, build a *de novo* SNP catalog and call genotypes. Following Mastretta□Yanes et al. (2015), we tested different sets of Stacks core parameters (ustack − m (2 to 6), −M (2 to 6) and −max_locus_stacks (2 to 6) and cstack −n (0 to 5) − Supplementary Material Figure S1 and S2), by varying one parameter at a time while holding the others at their default values. We then selected the set of parameters that minimized error rates between replicates (*n* = 28 pairs) and maximized the amount of data recovered. The optimal values were: -m (5), -M (2), -max_locus_stacks (3) and -n (2; error rates for each set of parameters are presented in Supplementary Material Figure S1 and S2). To produce the final data set, we ran Stacks with all parameters set to their optimal values. We performed the next filtering steps in *R*3.6.2* (R Core Team 2019) from the original VCF file obtained from stacks and by keeping only loci that were typed for at least 85% of samples and only samples that were typed for at least 80% of loci. We then used *dartR* (Gruber *et al*. 2018) and *Radiator* (Gosselin *et al*. 2017) packages in *R* and *PDGspider* (Lischer & Excoffier 2012) to convert SNPs VCF files to other formats (e.g. genlight; Jombart 2008).

### Population Genetic Structure, Genetic Distance, and Isolation-by-distance

We used complementary analyses to quantifying population genetic structure. Unless mentioned, all analyses were in *R*\*3.6.2 (R Core Team 2019). We first used Principal Component Analysis (PCA), considering the a priori defined wintering regions with the function *glPCA* using the package *Adegenet* *2.1.1 (Jombart 2008). Next, to compute pairwise *F*_ST_ among regions according to Weir and Cockerham (1984) we used the *gl.fst.pop* in the *dartR* package (Gruber et al. 2018; See the Supplementary Material Appendix). Also, we performed a Discriminant Analysis of Principal Components (DAPC; Jombart et al. 2010; also implemented in the package *Adegenet* (Jombart 2008)) to infer the number of genetic clusters that best fit our data. In addition, we used a Maximum Likelihood approach using ADMIXTURE*1.3 software (Alexander *et al*. 2009) to estimate the number of populations underlying the genetic dataset based on individual relatedness without our *a priori* wintering regions. Finally, we tested for Isolation-by-distance at the individual level by investigating the between the natural logarithm of Euclidean distance and relatedness (i.e. *beta*, according to Weir and Goudet 2017) distance matrices between each pair of individuals. An extended description of the methods is provided in Supplementary Material Appendix.

### Descriptive Genetic Diversity

We calculated the observed heterozygosity (*Ho*) and the expected heterozygosity (*He*) for each locus using the function *gl.Ho* and *gl.Hs* in the *dartR* (Gruber *et al*. 2018) package in *R*\*3.6.2 (R Core Team 2019). We also computed *Ho* and *Hs* for each wintering area. To test whether a significant difference in *Hs* among wintering areas can exist, we performed a Linear Mixed Model (LMM) with wintering area as the fixed effect and the loci ID as a random effect. We verified linearity assumptions of model residuals, and we then computed a post-hoc Tukey test to contrast wintering areas.

### Current Effective Population Size (*Ne*)

We used NeEstimator*2.1 (Do *et al*. 2014) to estimate the current effective population size (*Ne*) of Snowy Owls in North America. We used the linkage disequilibrium (LD) method with a random mating model. Following Waples and Do (2010), we excluded rare alleles (i.e., those SNP with a minor allele frequency ≤ 0.05) to avoid estimation bias.

### Population Trajectory Reconstruction

We reconstructed the population history of Snowy Owls using the Stairway Plot 2 method (Liu and Fu 2020). The Stairway Plot infer detailed population demographic history using the site frequency spectrum (SFS: Liu and Fu 2015) from DNA sequence data. This method uses a flexible, multi-epoch model as used in the Skyline Plot method (Strimmer and Pybus 2001, Navascués et al. 2017), and based on the expected composite likelihood of a given SNP frequency spectrum (SFS). It provides information on the history of population effective size (*Ne*) over time until the recent past. We estimated the folded SFS using the VCF file and a Python script, *EasySFS* (https://github.com/isaacovercast/easySFS; Covercast 2017) and included the total number of observed nucleic sites (both polymorphic and monomorphic sites). The “two-epoch” model, with 67% of sites for training and 200 bootstraps was used as recommended. We tested four different numbers of breaking points (i.e. to define the boundaries of each epoch; Liu and Fu 2020) as described in Liu and Fu (2015), i.e. 74, 149, 223, and 298. We assumed a mutation rate of 1.9 × 10^−9^ which is a recent estimate for birds in general (Zhang et al. 2014) and a generation time of 7.93 years (a generally accepted value; BirdLife International 2020). However, because we have no information on the mutation rate in Snowy owls, and because there is also some uncertainty over generation times in the literature (e.g. 4.7 years, Heggøy et al. 2017; 7.93, BirdLife International 2020), we ran sensitivity analyses using various mutation rates retrieved from the literature on birds (i.e. *μ*= 4.6×10^−9^, Smeds et al. 2016); *μ*=1.9 × 10^−9^; Nam et al. 2010, Zhang et al. 2014), and ranges of generation times. We also added the major past glaciation events to the resulting Stairway plot (Clark *et al*. 2009, 2012, Mann *et al*. 2009).

## RESULTS

### SNPs Genotyping

After the *de novo* SNP calling and filtering procedure, the data set encompassed 13,793 SNPs distributed over 5,987 loci. We only kept the SNPs that had less than 15% of missing data, i.e., that were typed for at least 85% of birds. Individuals scored on average 5,767 SNPs ± 198 (SD) [4,826; 5,921], resulting in 3.7% missing data in the genotype matrix. Error rates estimated on the 13,793 SNPs obtained from replicated samples were 0.004 +/- 0.003 and dropped to 0.003 +/- 0.002 when considering single SNP per loci (*n*=5,987 SNPs). For subsequent analyses, we only considered a single SNP per locus, for a final dataset consisting of 5,987 SNPs genotyped on 150 Snowy Owl individuals.

### Population Genetic Structure, Genetic Distance, and Isolation-by-distance

Results from the PCA (Supplementary Material Figure S3), the DACP (Supplementary Material Figure S4), and from ADMIXTURE (Supplementary Material Figure S5–S6) were congruent and all revealed a single genetic cluster (*K*=1). When Admixture was conducted separately on each dataset of yearling owls versus the adult owls, both also revealed a single genetic cluster (*K*=1). Pairwise *F*_ST_ between wintering areas averaged 7e^−5^ ± 3e^−4^ (Figure 1). However, pairwise *F*_ST_ analysis on adults from W and E, the two most distant wintering areas, revealed a small, but significant genetic differentiation (*F*_ST_ = 0.002; *p* = 0.017; Figure 1 and Supplementary Table S2), which also held for the adult females between W and E (*F*_ST_ = 0.002; p = 0.006; Supplementary Table S2) and the adult females between W and GL (*F*_ST_ = 0.001; *p* = 0.042; Supplementary Table S2). There was also a low genetic differentiation between males from W and C (*F*_ST_ = 0.002; *p*=0.042; Supplementary Table S2). Global *F*_IS_ was 0.0332 ± 0.2431. Mean *F*_IS_ were low to moderate (but all significantly different from zero; all Wilcoxon signed rank tests *p*-values < 0.001) and were similar within wintering areas: W= 0.037 ± 0.19; C= 0.038 ± 0.21; GL = 0.048 ± 0.22 and E = 0.040 ± 0.20.

On average, genetic relatedness was 0.017 ± 0.014 (Figure 2), and the maximum value between two individuals was 0.093, one sampled in the E wintering area (New York) and one in the GL area (Michigan; 2,763 km apart). The minimum value between the two individuals was 1.98 × 10^−6^, one from the E (New York) and one from the GL areas (Wisconsin; 2,457 km apart). Genetic distance was similar within and between wintering areas (Figure 2) and between adults and first year (Figure S7). We did not find any pattern of Isolation-by-distance (at a scale of 4,650 km), i) at the individual level (i.e. relatedness vs ln-transformed geographic distance): Mantel’s *r* = 0.005, *p* = 0.39 and LMM slope=-8.87 e^−5^, 95% CI= −1.86e^−4^; 9.30e^−6^, and ii) at the population level (i.e. *F_ST_ /* (1-*F*_ST_) vs ln-transformed geographic distance): Mantel’s *r* = −0.258, *p* = 0.70 and LMM; slope=-3.69 e^−4^, 95% CI= −6.8 e^−4^; 4.75e^−5^).

**Figure 2.**
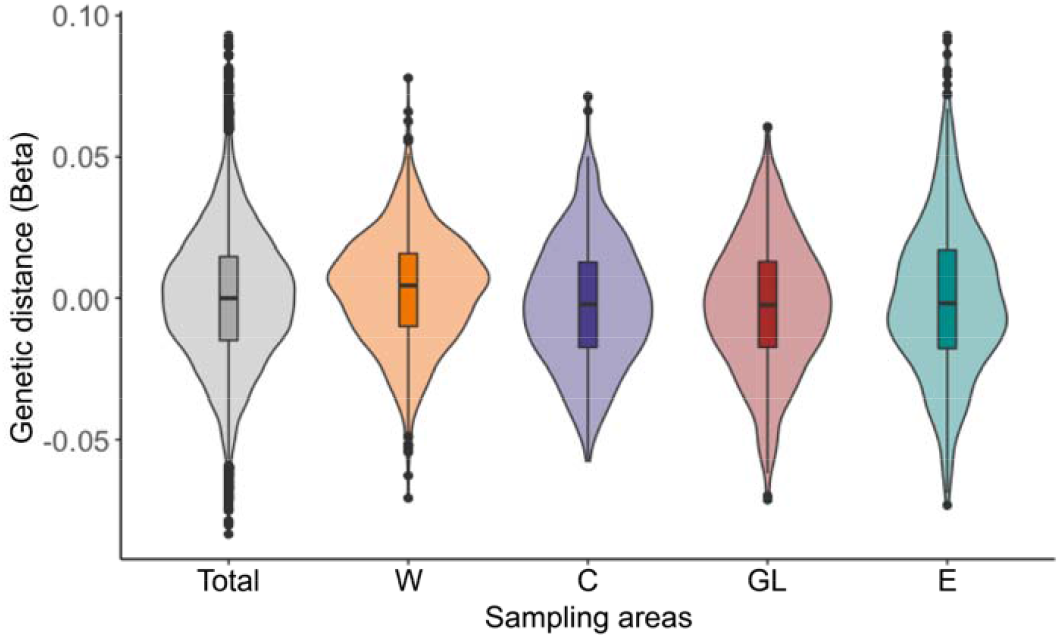
Violin plot of genetic distance [beta estimation of relatedness between every pair of individuals] for all individual Snowy Owls pooled (Total) and for each wintering area (W-West, C-Central, GL-Great Lakes, E-East).

### Descriptive Genetic Diversity

At the locus scale, the average observed heterozygosity (*Ho*) per locus was 0.142 ± 0.143 (SD) and the expected heterozygosity per locus (*He*) was 0.148 ± 0.140. At the wintering area scale, we found a significant decrease of gene diversity from West to East (*Hs*; slope= −1.847e^−2^, SE= 5.664e^−3^, 95% CI= −0.0296, −0.00737; Figure 2).

### Curent Effective Population Size (*Ne*)

Using the LD method with a lowest allele frequency set at 0.05, the current effective population size (*Ne*) for the North American population of Snowy Owls was estimated to be 15,792 individuals (95% CI 10,850-28,950). Excluding singleton alleles only, the *Ne* value was 15,401 (95% CI 12,782-19,366).

### Population Trajectory Reconstruction

The Stairway plot approach showed that the coalescence *Ne* of Snowy Owls in North America declined steadily in recent times (Figure 3). There was an expansion of *Ne* around 100,000 years ago, followed by a long period of stability. Coalescence *Ne* then started to decline around 6,000 years ago, which coincides with an acceleration of deglaciation in North America between 6,000 and 8,000 years ago. This is also approximately timed with the beginning of the Holocene period, which was characterized by global temperature increases. During the Little Ice Age period (*ca* 1,400-1,700 AD), the rate of decline in coalescence *Ne* apparently slowed down. The decline started to speed up again around 200 years ago, approximately at the onset of the worldwide retreat of glaciers and the acceleration of air temperature warming (Figure 3). We cannot infer population changes for the last 100 years because of the uncertainties in the final steps of the Stairway Plot method (Liu and Fu 2015, 2020). The sensitivity analysis showed that despite using different values for mutation rate or generation time, all demography simulations resulted in a declining trend (Supplementary Material Figure S8).

**Figure 3.**
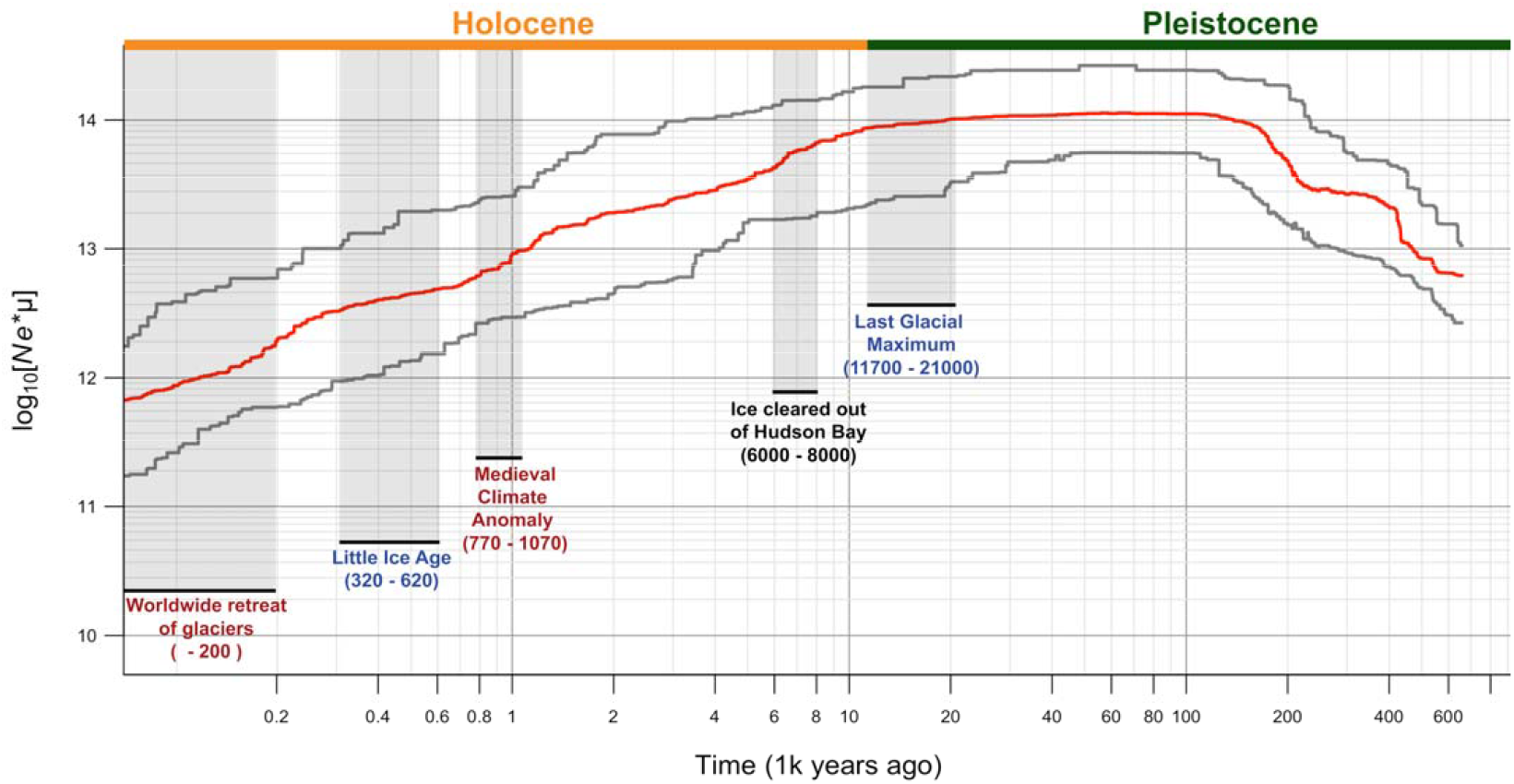
Reconstruction of the effective population size of the snowy owl population (*Ne*) from 600 Ka until 100 years ago in North America, assuming mutation rate of 1.9 × 10^−9^ and a generation time of 7.93 based on a Stairway Plot 2 (Liu and Fu 2020). The method precludes reliable estimates for the most recent 100 years. Grey lines indicate the 95% confidence interval. Light grey areas represent glaciation events according to Clark et al. (2009, 2012); Mann et al. (2009). Events in red are characterized by higher temperature and events in blue are characterized by lower temperatures.

## DISCUSSION

### Continent-scale Genetic Homogeneity

Based on the information retrieved from Snowy Owls on the wintering grounds, our results all indicate that populations at the continental scale are genetically homogenous in North America. Two non-mutually exclusive hypotheses could explain the absence of genetic differentiation on a 4,500 km scale: breeding dispersal or juvenile (natal) dispersal. Irrespective of their wintering areas, if owls disperse widely during the pre-breeding period or between successive reproduction events (i.e., breeding dispersal, Clobert et al. 2012) individuals from different wintering areas may come into contact, mate, and breed together in the high Arctic. There is some evidence that Snowy Owls have low breeding site fidelity and that long distances may separate breeding sites in consecutive years (Fuller et al. 2003, Therrien et al. 2014, Doyle et al. 2017, Holt et al. 2020). Thus, mating is likely panmictic and not related to the winter origin of birds.

A second possibility is that high juvenile dispersal from the site of birth to the site of reproduction is driving the genetic mixing. Natal dispersal distance is typically much larger than breeding dispersal in many birds (Clobert et al. 2012) and probably plays a role in genetic intermixing in other owl species such as the Northern Spotted Owl; *Strix occidentalis caurina*; (Miller *et al*. 2018), but nothing is known about natal dispersal in Snowy Owls. This hypothesis could explain the absence of genetic differentiation if juveniles are dispersing among wintering areas during the first years of life, i.e. before their first reproduction. Alternatively, genetic homogenization could occur as a result of periodic winter irruptions when, after a summer with high productivity linked to abundant lemming prey in the Arctic, large numbers of juveniles travel farther south than average during their first winter (Kerlinger & Lein 1988, Potapov & Sale 2012, Holt *et al*. 2020). High gene flow is probably facilitated by the high mobility and nomadic behavior of the species, allowing high breeding and natal dispersal or a combination of both (Yannic et al. 2016).

The pairwise *F_ST_* values among winter areas of Snowy Owls (ranging −0.0003 to 0.0004) are low compared to the range of values based on SNPs from other raptor species. Gousy-Leblanc *et al*. (2021) recently reviewed genetic differentiation within raptors and reported values for four species based on data from breeding grounds at smaller scales than our current study and with different ecology than Snowy Owls (*i.e*., fully sedentary, or partially migratory species): Great Gray Owl *Strix nebulosa F_ST_* = 0.03-0.15 (Mendelsohn *et al*. 2020), Prairie Falcon *Falco mexicanus F_ST_* = 0.01-0.03 (Doyle *et al*. 2018), Bald Eagle *Haliaeetus leucocephalu; F_ST_* = 0.037-0.203 (Judkins *et al*. 2019), and Northern Goshawk *Accipiter gentilis; F_ST_* = 0-0.093 (Geraldes *et al*. 2019). All estimates were at least an order of magnitude higher than those we found in Snowy Owls, and none of these species had movements as nomadic as those observed in Snowy Owls.

Conversely, the Gyrfalcon (*Falco rusticolus*), an Arctic raptor which inhabits similar habitats as the Snowy Owl, shows genetic differentiation between the continental populations of Greenland and Iceland, despite the high dispersal capability of the species (Johnson *et al*. 2007). For the Eurasian Arctic Peregrine Falcon (*Falco peregrinus*), four distinct genetic groups were detected at the continental scale, which supposedly arose from the use of distinct population-specific migration routes, and fidelity to breeding and non-breeding areas (Gu *et al*. 2021). At such a large scale and for species with a high dispersal capacity, various factors like fidelity to breeding or non-breeding areas, low breeding and natal dispersal, the use of distinct migration routes, and the interplay between those factors could impede gene flow among populations, promoting genetic differentiation.

### Current Genetic Diversity and Effective Population Size (*Ne*)

For Snowy Owls, we found a slight statistically significant genetic diversity between the most distant regions, i.e., the Western and Eastern regions in North America. This could be explained by the potential inclusion (i.e. via migration) of individuals in the “Western” category which originated from Alaska or Russia but whose dispersal was not far enough to reach the Eastern population (Fuller *et al*. 2003, Holt *et al*. 2020). Such heterogeneous migration could slightly influence allele frequencies across the circumpolar distribution, while not greatly impacting the genetic differentiation overall. Indeed, we observed that the highest *Hs* occured in the West. The global population should nonetheless be considered as one population unit genetically.

The average expected heterozygosity (0.148 ± 0.140 SD) at the locus scale for Snowy Owls is slightly lower compared to the *He* values computed with SNPs for three other species of raptors. Two species that were partial migrants displayed genetic differentiation at large scale, *He* = 0.293 in the Bald Eagle, *H. leucocephalus* (Judkins *et al*. 2019) and *He* = 0.340 in the Golden Eagle *Aquila chrysaetos* (Doyle *et al*. 2016). One obligate migrant species, the Prairie Falcon *Falco mexicanus*, showed no population genetic differentiation at large scale *He* = 0.332 (Doyle *et al*. 2018). However, *He* in Snowy Owls is quite similar to the Burrowing Owl *Athene cunicularia*, another nomadic facultative migrant species (*He* = 0.112) that, exhibited population genetic differentiation at low scale (Mueller et al. 2018).

Although Snowy Owls appear to have lower gene diversity than other raptors, the estimated current effective population size of the species in North America (*Ne* = 15,792 individuals) is large enough to preserve the evolutionary potential of the population and allow it to persist and adapt in a changing environment as suggested by population modelling (see Kamath et al. (2015) and Hoban et al. 2020). In general, a minimum *Ne* of 100 individuals is recommended to prevent loss of genetic diversity by genetic drift and a *Ne* of 1,000 individuals to maintain long-term evolutionary potential (Frankham *et al*. 2014). In a conservation context, *Ne/Nc* where *Nc* is the number of individuals counted in a census population, is important for disentangling the relative effects of population size and genetic factors on the persistence of species (Frankham 1995, Palstra & Ruzzante 2008, Ferchaud *et al*. 2016, Waples 2016). Considering our estimated *Ne* of 15,792 individuals (95% CI between 10,850-28,950) and a *Nc* of about 30,000 individuals in North America (Rosenberg *et al*. 2016, BirdLife International 2020), we can calculate a *Ne/Nc* ratio of about 0.57 [range 0.36 - 0.96]. Estimates of *Nc* for Snowy Owls, are extremely uncertain and may be as low as 7,000-8,000 pairs in North America (Potapov & Sale 2012, BirdLife International 2020). *Ne/Nc* ratio values derived from these lower *Nc* estimates would then be around 1.05 [0.77 - 1.81]. All these *Ne/Nc* values are nonetheless higher than those in many other avian taxa (Frankham 1995) so our results suggest that the North American population as a whole does not face an imminent risk of inbreeding depression or genetic impoverishment.

### Past Population Trajectory

Demographic reconstruction suggests that the population size of Snowy Owls declined over the last few thousand years, starting around the Last Glacial Maximum (LGM). A recent reconstruction of demographic history in the Peregrine Falcon (*Falco peregrinus*), another Arctic raptor, resulted in a similar population size trajectory as the Snowy Owl (i.e. expansion during the LGM and a steady decline since the beginning of the Holocene; Gu et al. 2021). Perhaps an increase in the total area of tundra habitat may explain the population expansion during the LGM and the subsequent decline in the population occurred when the tundra began to contract northward (Gu *et al*. 2021). Similarly, the population size of Snowy Owls may be closely matched to the amount of Arctic tundra because the species breeds exclusively in that habitat. An implication is that large and long-term environmental changes over the last few thousand years, and in particular to temperature increases since the LGM (Clark et al. 2009) may greatly affect the population size of Snowy Owls. Indeed, population reconstructions showed that temperature increases reduced the effective population size of three penguin genera (*Eudyptes, Pygoscelis*, and *Aptenodytes*) in Antarctica (Cole *et al*. 2019), highlighting ecosystem-wide responses to climate changes in the Antarctic ocean in the past. Similarly, the climate warming that occurred after the LGM in the Northern Hemisphere reduced effective population sizes, genetic diversity, and/ or census counts of other Arctic species like the polar bear *Ursus maritimus* (Miller *et al*. 2012) and the Arctic fox *Vulpes lagopus* (Larsson *et al*. 2019). Global warming has also been implicated as a major factor in the mass extinction of Late Quaternary megafauna in the Northern Hemisphere (Lorenzen et al. 2011, Lord et al. 2020, Stewart et al. 2021; but see Sandom et al. 2014).

Globally, the warming period of the Holocene has been an important influence on population size and/or range distribution of small mammals including lemmings (Prost *et al*. 2010, 2013, Lanier *et al*. 2015, Fedorov *et al*. 2020) which are the main prey of breeding Snowy Owls. Thus, climate change has the potential to influence entire ecosystems in the Arctic by disrupting the main trophic links between species and Snowy Owls may have declined since the mid-Holocene both as direct result of warmer temperatures (e.g. physiological stress) and/or due to the indirect effect of the climate-induced population reductions and geographic range contractions of its main prey, i.e., the small mammals in the Arctic.

## Conclusion

Our findings should help further define some research priorities for Snowy Owls identified by Holt et al. (2020). First, the lack of significant genetic differentiation over the span of a continent spanning > 4,500 km does not support the hypothesis that well-separated migration routes, i.e., north-south corridors to distinct winter grounds are serving to isolate distinct breeding populations of Snowy Owls. Second, and potentially most concerning, our results showed that Snowy Owls experienced a continuous decline in numbers since the end of the Last Glacial Maximum (18-21k years ago) in North America. Although our estimate of current effective population size does not suggest that Snowy Owls are at risk for imminent genetic problems, the finding that the owls were highly sensitive to global changes in the past implies that the population size may decline more precipitously in the future if Arctic warming accelerates, as predicted by many climate changes models.

## ACKNOWLEDGEMENTS

This work was supported by the Canada Research Chair Program, Polar Knowledge Canada, NSERC, and the Canadian Foundation for Innovation (N.L.). M.G.L. benefited from a NSERC and a FRQNT graduate grant. We thank C. Miquel as well as the AnaBM (USMB) and AEEM (UGA) laboratory facilities for their help and support during the laboratory work. We thank everyone that contributed samples, including D. Brinker, D. Genesky, G. Jacobs, T. McDonald, S. Weidensaul, A. Robillard as well as Project SNOWstorm members and supporters. In BC, all of the following organization contributed snowy owl carcasses: BC Ministry of Agriculture, Food and Fisheries, BC Ministry of Environment and Climate Change staff from multiple regions, BC Wildlife Park, Hillspring Wildlife Rehabilitation Center, Mars Wildlife Rescue Centre, North Island Wildlife Recovery Assoc., Northern Lights Wildlife Shelter, Orphaned Wildlife Rehabilitation Soc., Northern Wildlife Rescue, South Okanagan Rehabilitation Centre for Owls, University of Northern British Columbia, Vancouver Airport Authority, and Wildlife Rescue Assoc; in the Maritimes: P-Y Daoust, D. Weeks, F. de Biep. For helping to trap owls in Saskatchewan, we thank D. Zazelenchuk and M. Stoffel. This work benefited from access to the Biogenouest genomic platform at Station Biologique de Roscoff and we are grateful to the Roscoff Bioinformatics platform ABiMS for providing storage and computing resources. We thank E. Milot and A.-M. Dion-Coté for the Ms review.

## Ethics statement

All live captures followed Canadian Animal Care Committees guidelines.

## Author contributions

Conceived the idea, design, experiment: N.L., M.G.-L., G. Y., and J-F. T. Data collection: M. G-L, J-F. T., I. S., L. W., J. T. E., V. B, G. G., K. L. W., and N.L. Performed the laboratory work: D. R., N. C-G-G, S. T. and N. T. Analyzed the data: M. G-L., G. Y., T.B., N.L., and J-F. T. Wrote first drafts: M.G-L, G.Y., N.L., J-F. Final version of the paper revised and agreed by all authors.

## Data depository

Data used in the article can be found on Figshare: https://figshare.com/s/f1f01dc772d2f8c8698c.

## SUPPLEMENTARY MATERIAL

### Population genetic structure, genetic distance and Isolation-by-distance

Unless mentioned, all analyses were in *R* (3.6.2.; R Core Team 2019). We first computed a Principal Component Analysis (PCA) with the function *glPCA* using the package *Adegenet*\*2.1.1 (Jombart 2008) using the four wintering regions (W, C, GL and E), retaining 100 PCA axes according to the eigenvalues plot. To infer the number of clusters of genetically related individuals to best fit our data, we used a Discriminant Analysis of Principal Components (DAPC; Jombart et al. 2010) also implemented in the package *Adegenet* (Jombart 2008). We followed the recommendation from Miller et al. (2020) to report the methods and results. Without prior information (*de novo*, unsupervised DAPC), we computed the function *find.cluster(*) to determine the optimal number of clusters. Precisely, we ran successive *K*-means clustering with an increase in the number of clusters (K=1-10). We used the smallest Bayesian information criterion (BIC) value associated with K for defining the number of clusters. We explored grouping individuals by sex or age (adult only), but we did not find any pattern. We also used Admixture v 1.3 (Alexander *et al*. 2009) to estimate the number of clusters in our dataset. We ran 10 replicates of Admixture (without any prior) allowing for the number of clusters (*K*) in the model to vary from 1 to 10. We generated random seeds for each run (beta < 0.0001). We chose the *K* value that minimized cross-validation error and hence best fit the data (Alexander *et al*. 2009). The output generated from 10 independent runs across each *K* was summarized and graphically represented using the main pipeline implemented in CLUMPAK with default options (Kopelman *et al*. 2015). We performed 10 independent runs of ADMIXTURE with adults and first years separated. Additionally, we computed pairwise *F*_ST_ among wintering areas (W vs. C vs. GL vs. E) with the *gl.fst.pop* in the *dartR* package (Gruber *et al*. 2018). We calculated pairwise *F*_ST_ by gender (male *vs*. female) and age group (adult *vs*. first year) among wintering areas. We computed global and within wintering regions *F*_IS_ using *gl.basics.stats* from *DartR* and tested if mean *F*_IS_ were significantly different from zero using Wilcoxon signed rank tests with continuity correction. We computed a *beta* estimator genetic distance matrix (i.e. a relatedness estimation matrix; according to Weir and Goudet 2017) between each pair of individuals with the function *snpgdsIndvBeta* in the package *SNPRelate* (Zheng *et al*. 2012). Generated values were homolog to the *F*_ST_ genetic distance (Weir and Cockerham 1984) measured between pairs of individuals instead of pairs of populations (Weir and Goudet 2017). A value of 0.5 is a first degree of relatedness (expected e.g., for full-sibs) and 0.25 a second degree (e.g. between half-sibs). We also performed a genetic distance matrix between all individuals located in the same wintering areas and a genetic distance matrix per age group (adult *vs*. first year). We calculated Geographic Euclidean distances between each pair of individuals using distance matrix in QGIS v 3.8. (QGIS Development team 2020). We assessed Isolation-by-distance (IBD; Wright 1943), the tendency for genetic similarity to reflect geographic proximity (Meirmans 2012), with two analyses. First, we computed a Mantel test (Mantel 1967) between the *beta* (i.e. genetic distance matrix) and the geographic matrices using the mantel function in the *Vegan* package (Oksanen *et al*. 2013) with 10,000 permutations to assess significance. Second, we also used a maximum-likelihood population-effects model (MLPE) in a linear mixed model (LMM), in which the covariate structure is specifically designed to account for the specific non-independency among column and row values in an n x n distance matrices (Van Strien *et al*. 2012). We fitted LMMs with *lme4* package (Bates *et al*. 2015). We verified linearity assumptions. We also ran the same analysis (Mantel and LMM) at the population scale, i.e. testing the relationship between the logarithm of Euclidian geographic [km] and the pairwise genetic distance *F*_ST_/(1-*F*_ST_) among all pairs of wintering regions following (Rousset 1997).

### TABLES

**Table S1:**
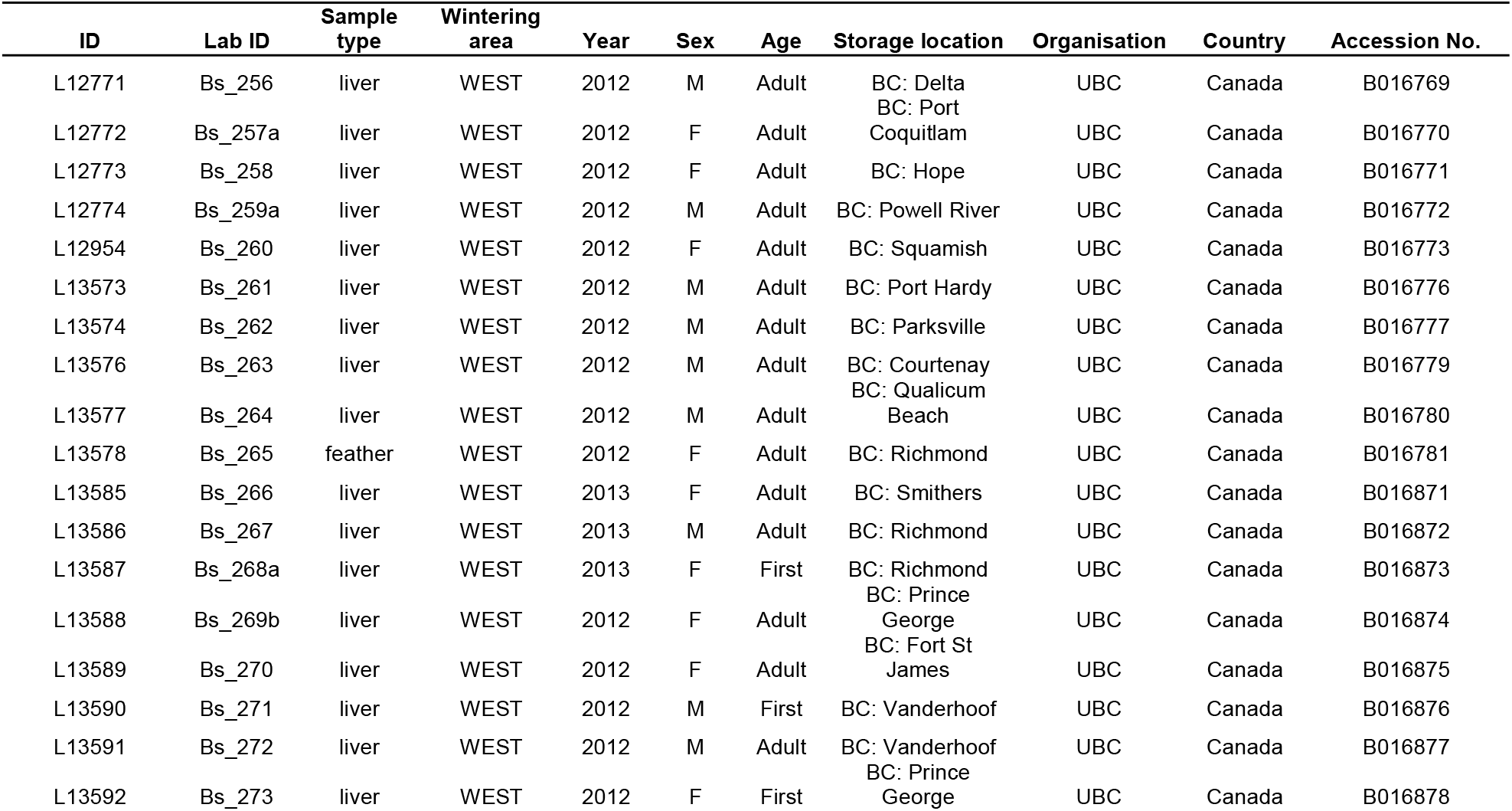
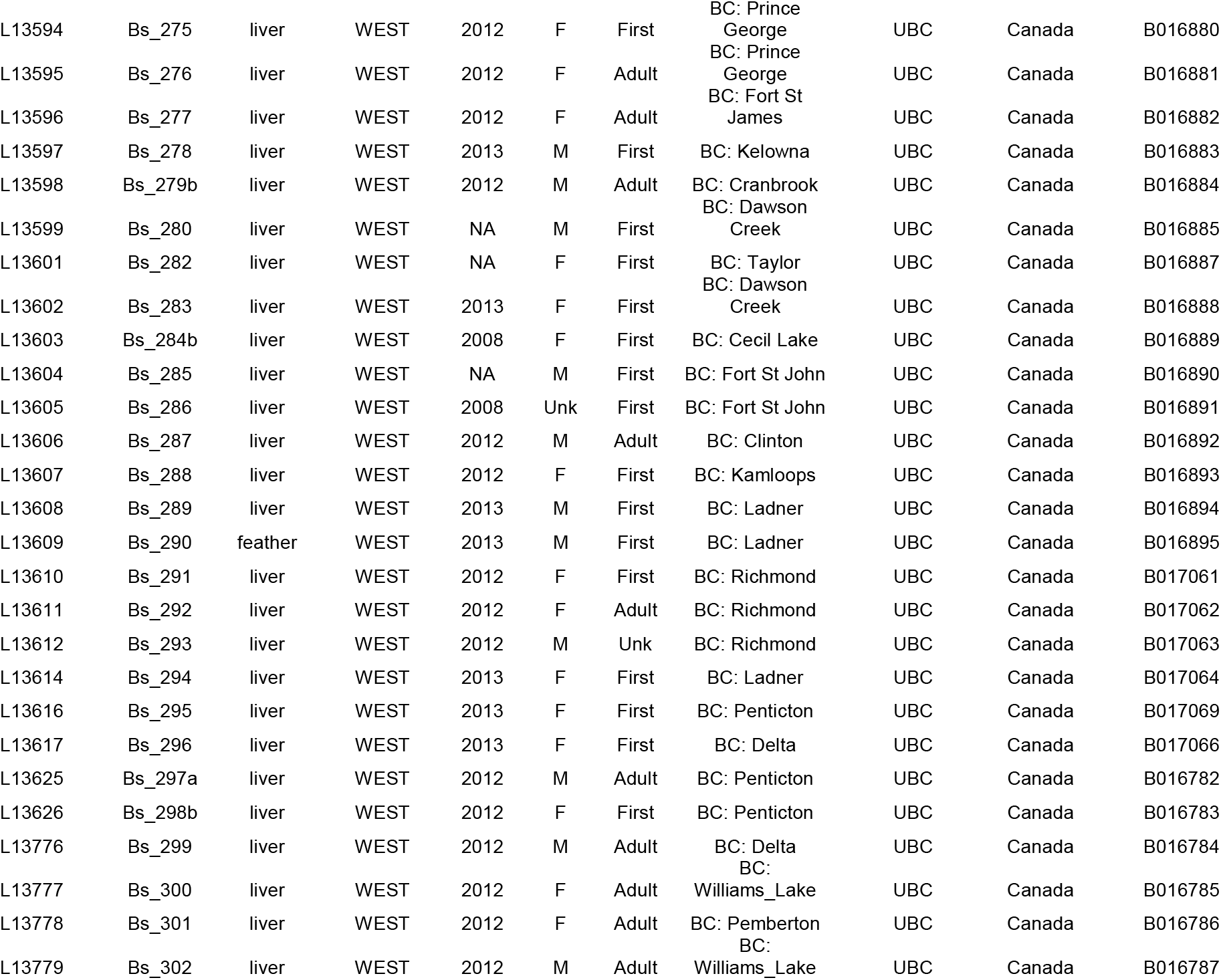
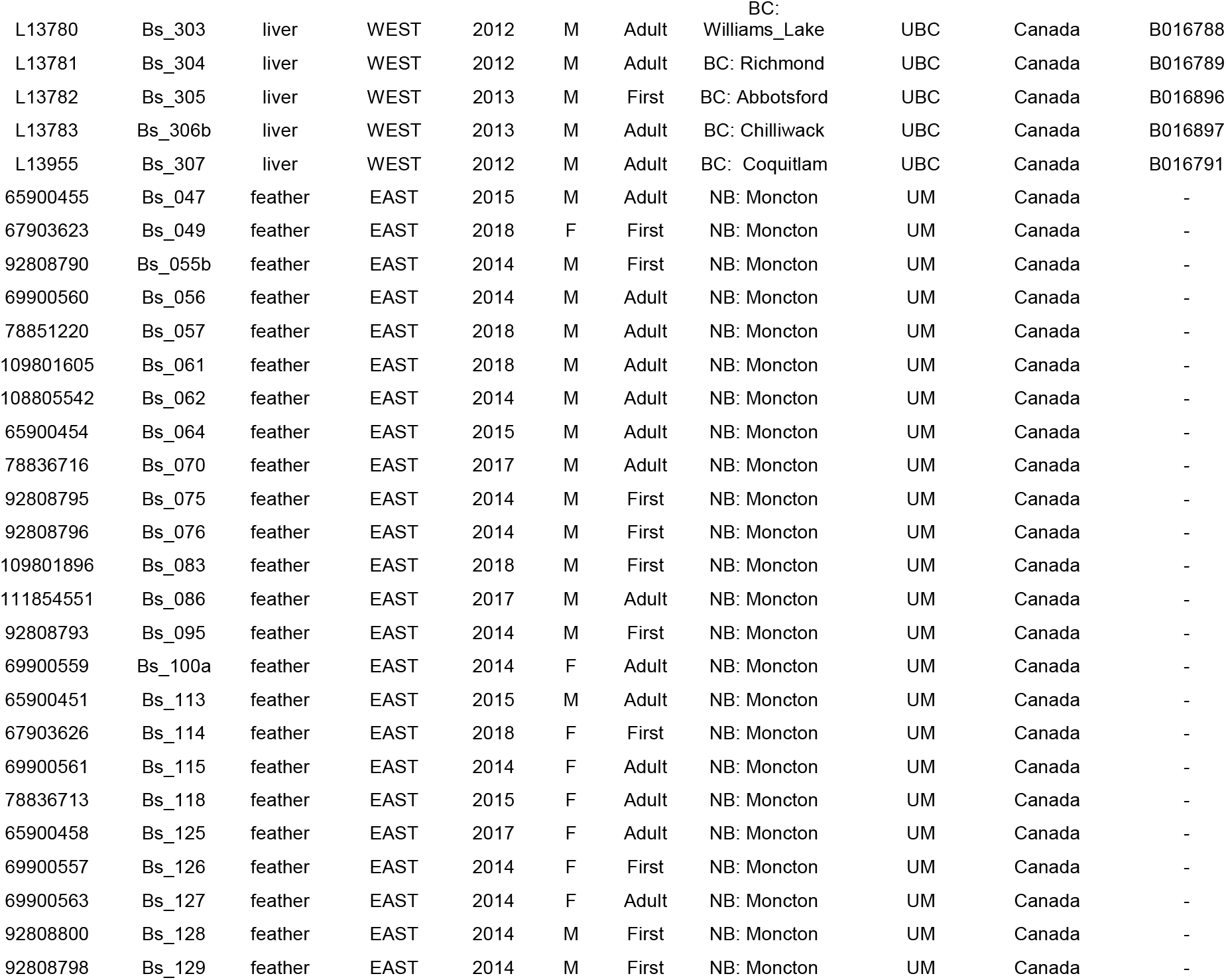
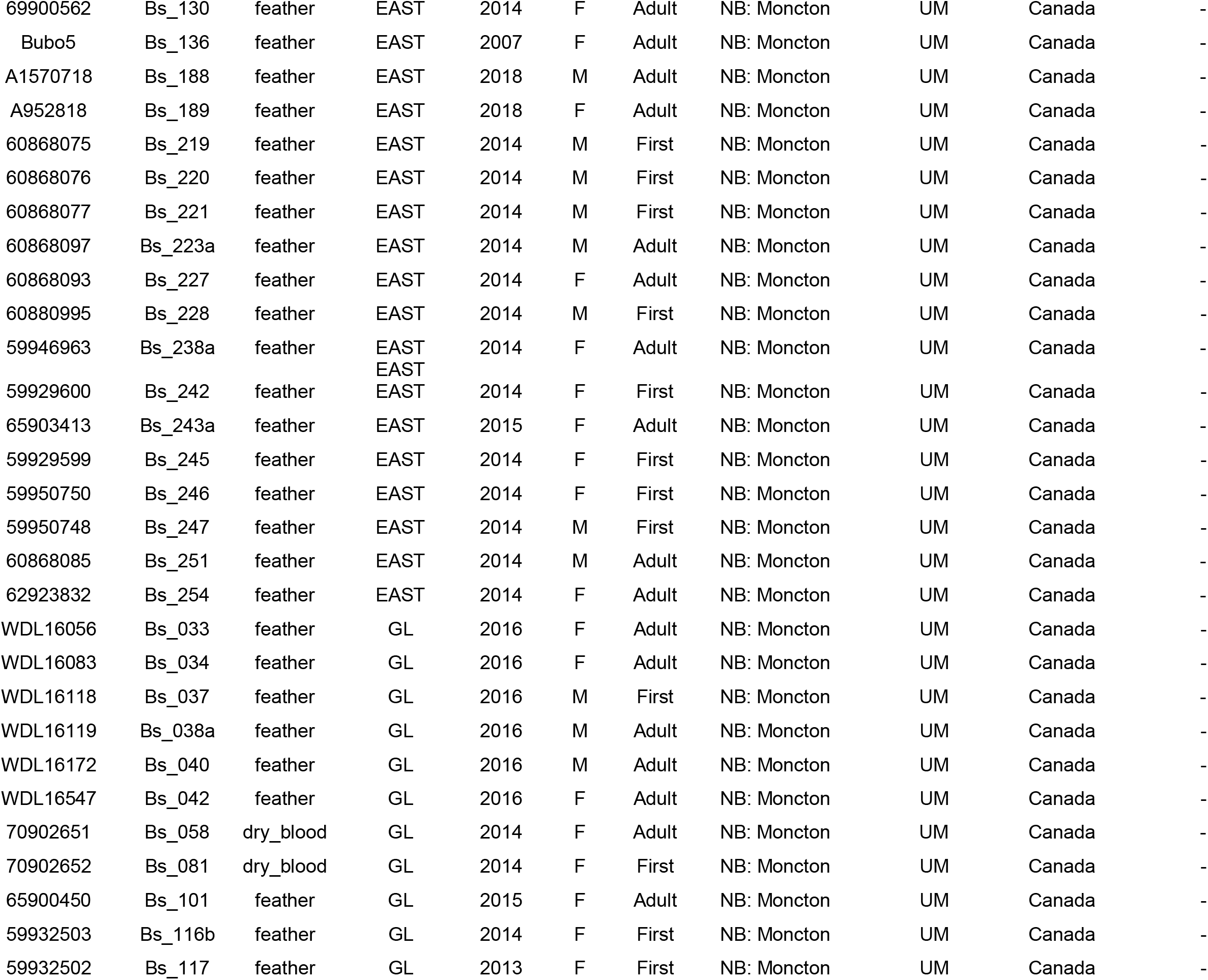
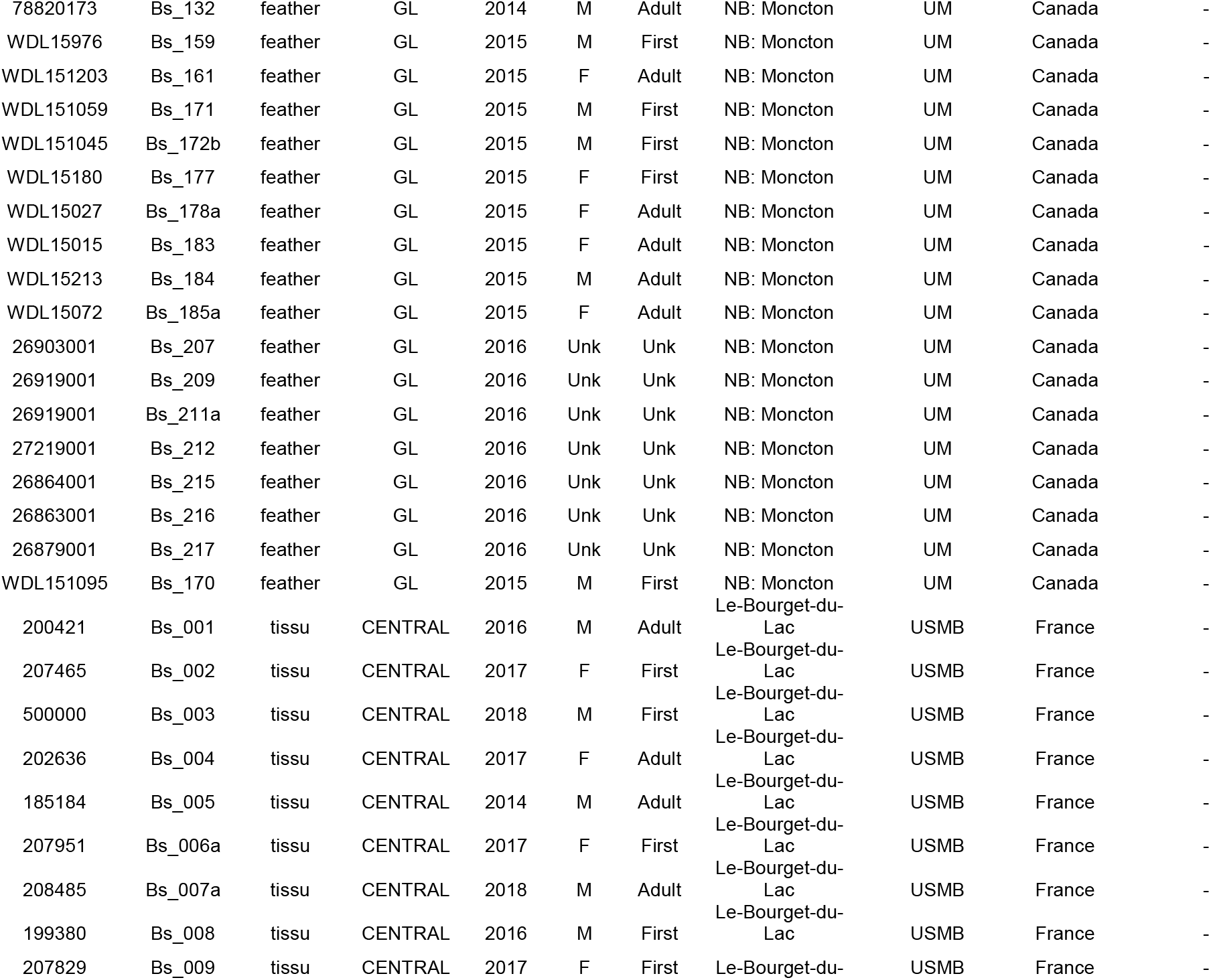
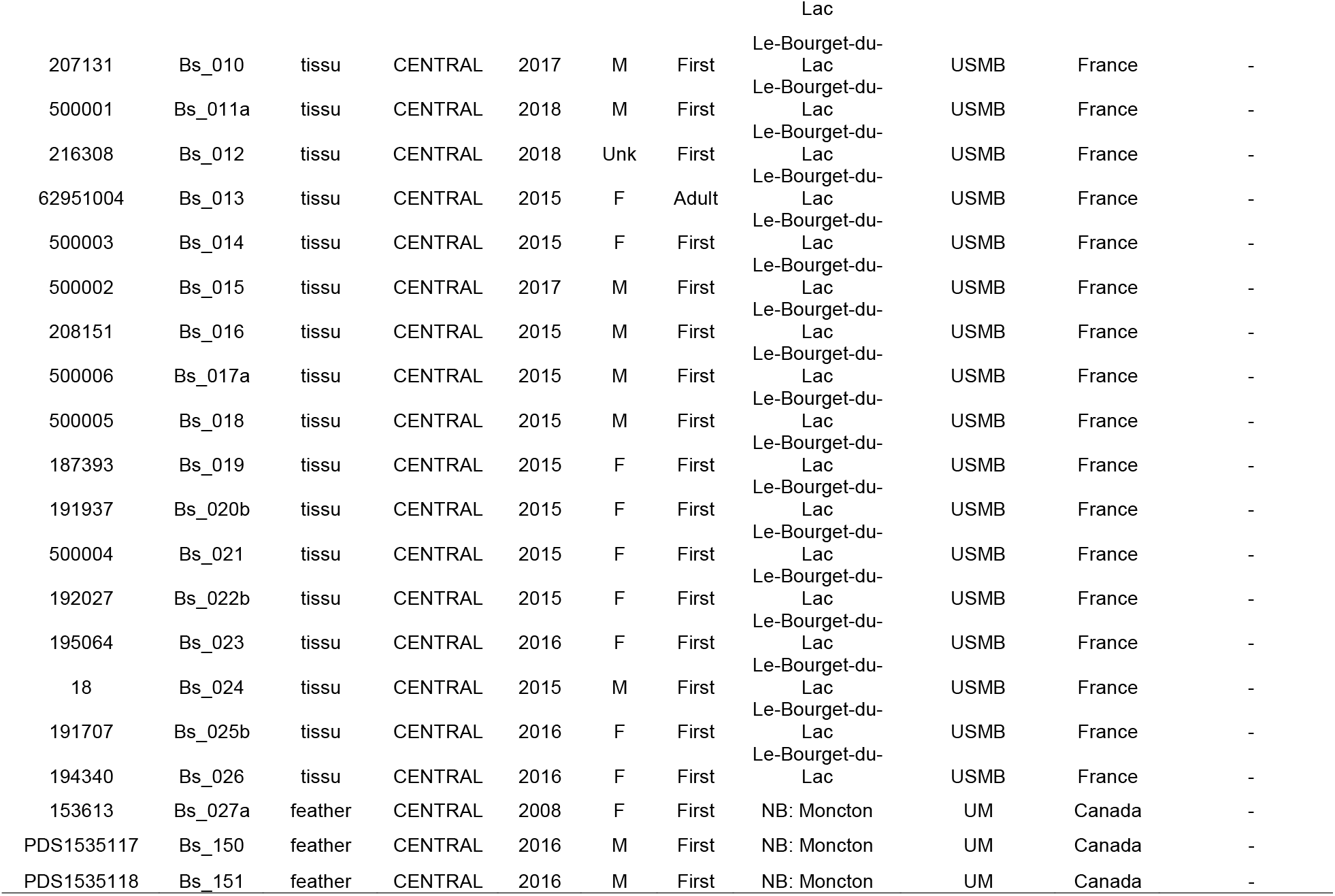
Sample information. **ID:** unique sample identification; **Lab ID:** identification used during laboratory work (UBC Beaty Biodiversity; UM Université de Moncton; USMB Université Savoie Mont-Blanc); **Sample type**: liver, tissue, feather or blood; **Wintering area:** West (British Columbia); Central (Saskatchewan); GL, Great lakes areas (Michigan, Wisconsin, Ontario, and Maryland); and East, i.e. all other individuals from the east coast of North America including the single Nunavut sample; **Year:** year of collection; **Sex:** sex of the individual; Age: Adult or first year (First); **Stored location:** actual location and long-term storage of the sample.

**Table S2:**
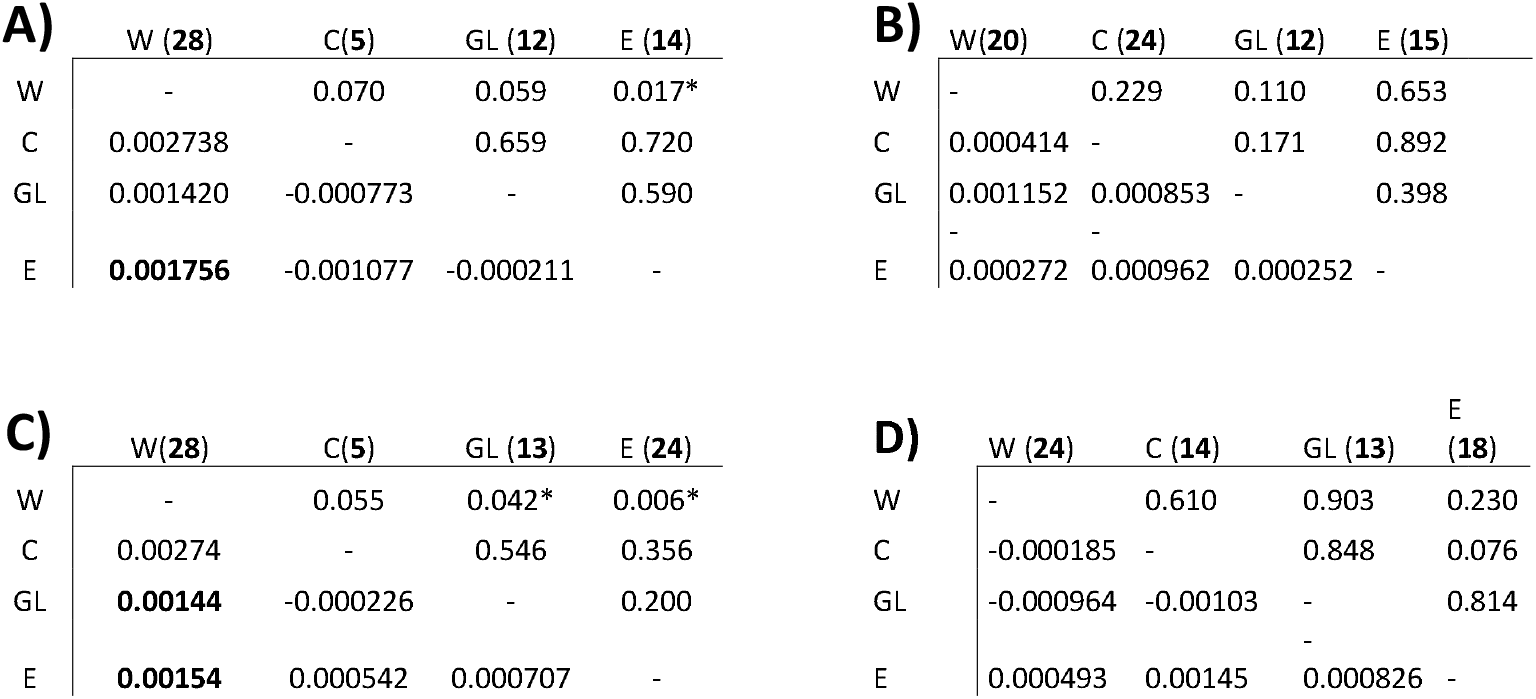
Pairwise *Fst* value (below diagonal) and p-value (above diagonal) between wintering areas (West (W), Central (C), Great lakes (GL), East (E) for A) adult individuals only B) first-year individuals only C) female only D) male only. Significant values are represented by a star and sample size per area is in bold. Negative values are equivalent to *F*_ST_ = 0.

### FIGURES

**Figure S1.**
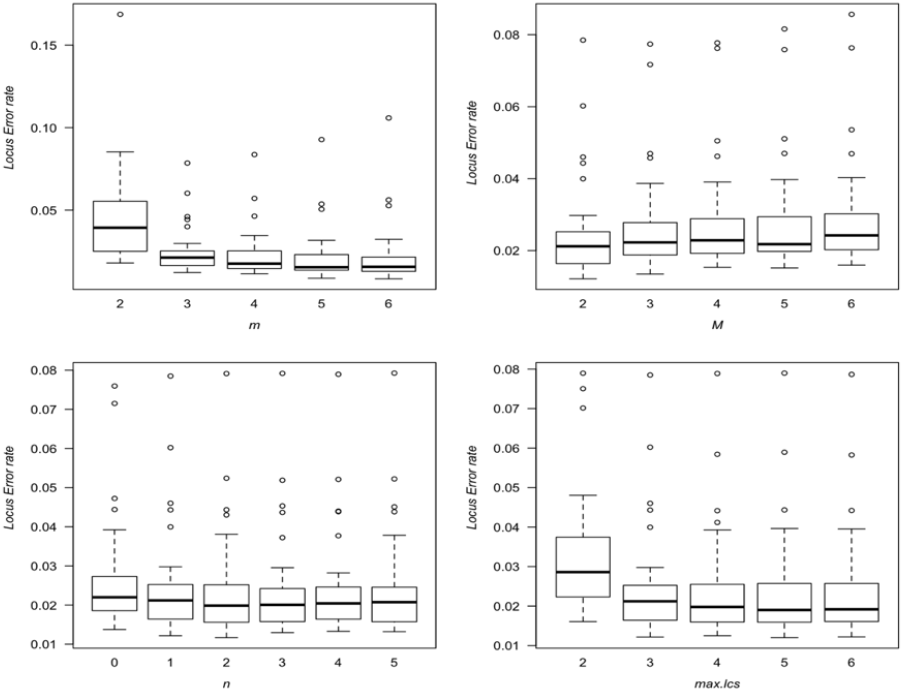
Effect of the variation of different *Stacks* core parameters locus error rated (proportion of missing loci in only one of the two replicates of a pair). Each box represents the result of a single *Stacks* run in which all parameters were set to their default values (m = 3, M = 2, N = M+2, n = 1, max_locus_stacks (mx.lcs) = 3, model = SNP) except for one that varied (indicated on the x-axis). See Mastretta-Yanes et al. (2015) for further discussion on parameters.

**Figure S2.**
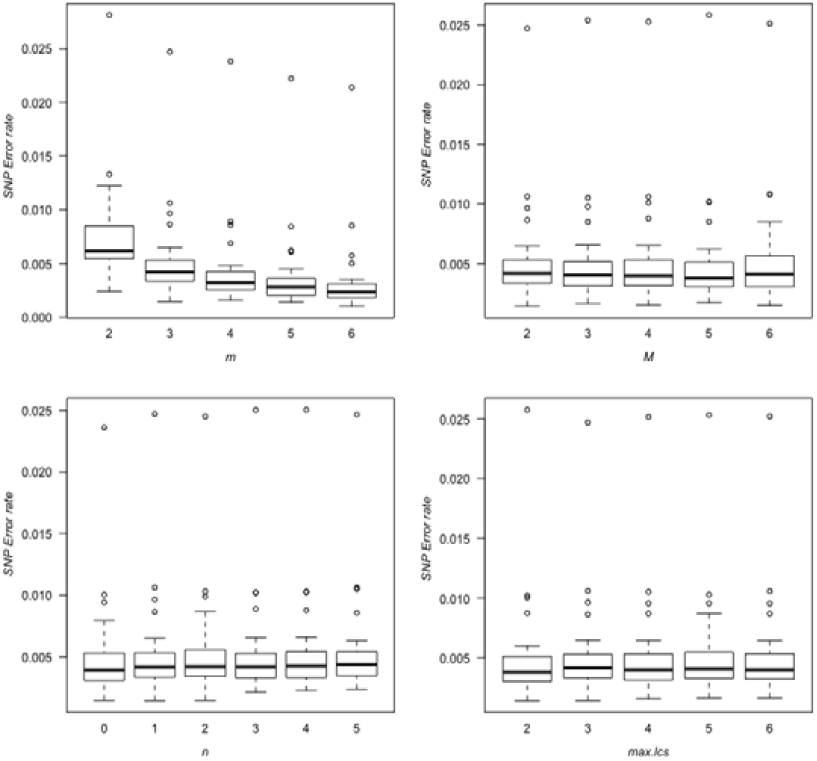
Variation of different *Stacks* core parameters on the SNP error rate (proportion of SNP mismatch within a replicate pair). Each box represents the result of a single *Stacks* run in which all parameters were set to their default values (m = 3, M = 2, N = M+2, n = 1, max_locus_stacks (mx.lcs) = 3, model = SNP) except for one that varied (indicated on the x-axis). See Mastretta-Yanes et al. (2015) for further discussion on parameters.

**Figure S3.**
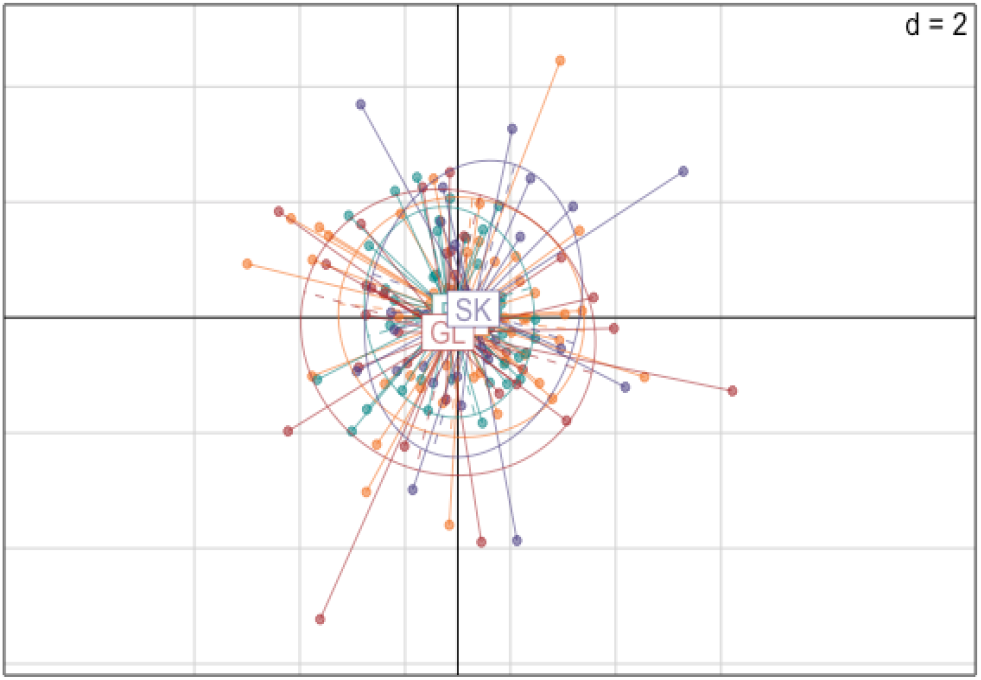
PCA of all the individuals according to their wintering areas. The number of retained PCs was 100. The number of dimensions (d) of the PCA is 2. Color codes: Orange=West (W, British Columbia, n=50), Purple=Central (SK, Saskatchewan, n=29), Red= Great lake (GL, n=29) and Green=East (E, n=42). The boxes with the letter are the centroid of each group.

**Figure S4.**
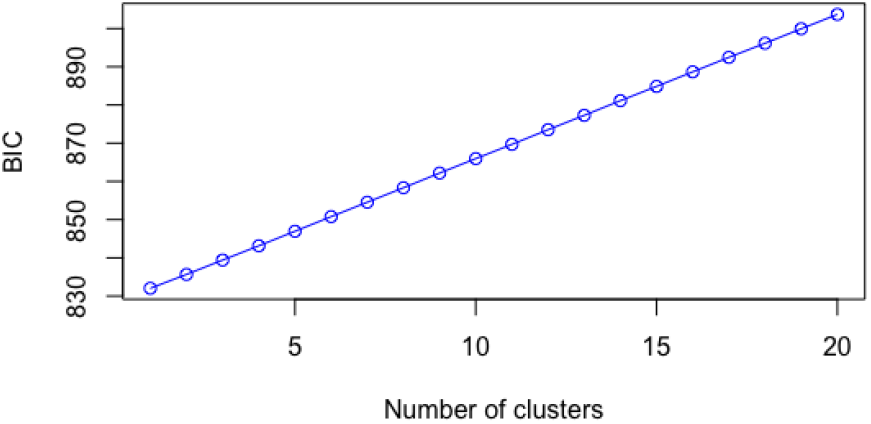
Relationship between the Bayesian information criterion (BIC) value and the number of clusters in the DAPC model.

**Figure S5.**
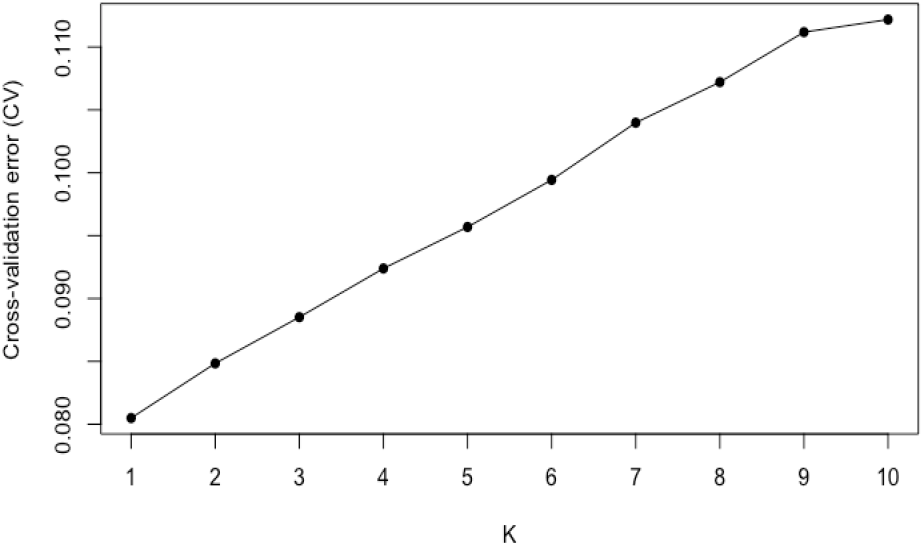
Cross-validation results of 10 runs (K=1-10) from the ADMIXTURE software. Selected value of the number of clusters (K) should be the lowest value (Alexander et al., 2009)

**Figure S6.**
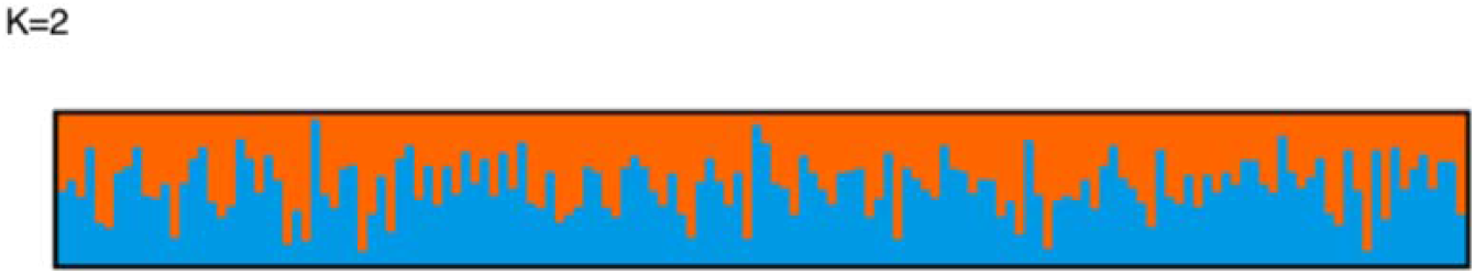
CLUMPAK results for *K*=2 from 10 runs of ADMIXTURE, without any prior allowing for the number of clusters (*K*) in the model to vary from 1 to 10. We generated random seeds for each run (beta < 0.0001).

**Figure S7.**
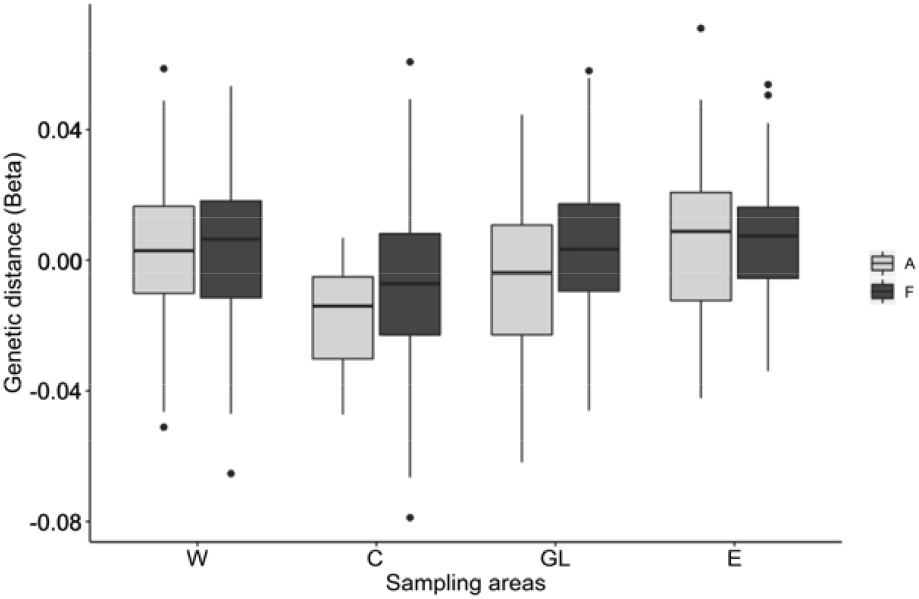
Boxplot of genetic distance (relatedness, beta index) of adults (dark grey) and first year (light grey) for each wintering area.

**Figure S8.**
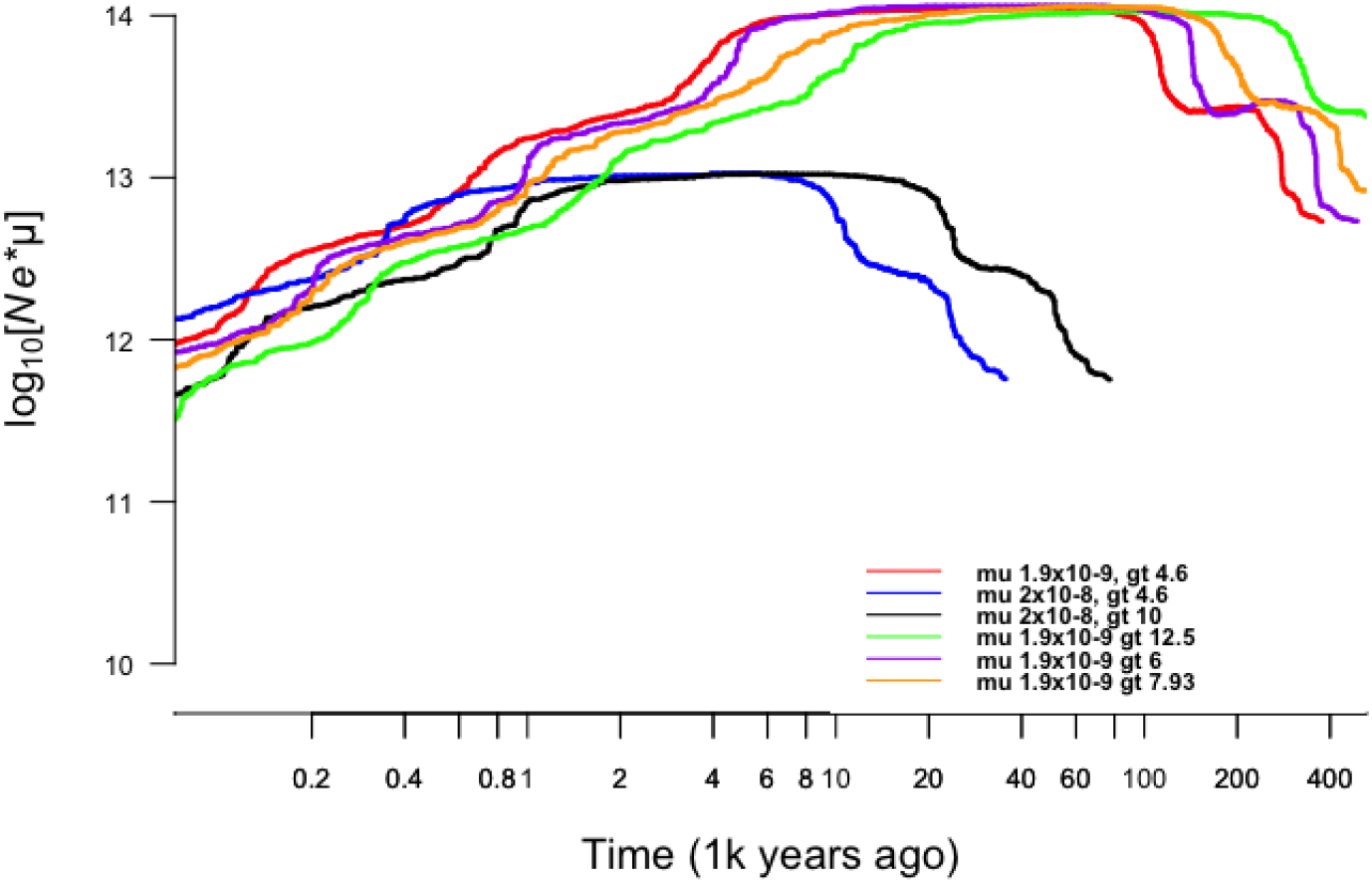
Stairway plot results of runs considering different mutation rates (μ: 2 × 10^−8^ and 1.9×10^−9^) or generation time (gt: 4.7, 6, 7.93, 10, and 12.5 years). For the sake of clarity, we did not provide the 95% confidence intervals in the figure (all very similar to the one presented in Fig. 3). The orange slope is the one that was selected in Figure 3.

